# Initiator AUGs are discriminated from elongator AUGs predominantly through mRNA accessibility in *C. crescentus*

**DOI:** 10.1101/2022.10.10.510831

**Authors:** Aishwarya Ghosh, Mohammed-Husain M. Bharmal, Amar M. Ghaleb, Jared M. Schrader

**Author notes:** These authors contributed equally to the work (Aishwarya Ghosh, Mohammed-Husain M. Bharmal). Center of Molecular Medicine and Genetics, Wayne State University, Detroit, MI, USA.

## Abstract

Translation initiation in bacteria is thought to occur upon base-pairing between the Shine-Dalgarno site in the mRNA and anti-Shine-Dalgarno site in the rRNA. However, in many bacterial species, such as *Caulobacter crescentus*, a minority of mRNAs have Shine-Dalgarno sites. To examine the functional importance of Shine-Dalgarno sites in *C. crescentus*, we analyzed the transcriptome and found more Shine-Dalgarno sites exist in the coding sequence than preceding start codons. To examine the function of Shine-Dalgarno sites in initiation we designed a series of mutants with altered ribosome accessibility and Shine-Dalgarno content in translation initiation regions (TIRs) and elongator AUG regions (EARs). A lack of mRNA structure content is required for initiation in TIRs, and when introduced into EARs, can stimulate initiation, suggesting that low mRNA structure content is a major feature required for initiation. SD sites appear to stimulate initiation in TIRs, which generally lack structure content, but SD sites only stimulate initiation in EARs if RNA secondary structures are destabilized. Taken together, this suggests that the difference in secondary structure between TIRs and EARs directs ribosomes to start codons where SD base pairing can tune the efficiency of initiation, but SDs in EARs do not stimulate initiation as they are blocked by stable secondary structures. This highlights the importance of studying translation initiation mechanisms in diverse bacterial species.

## Introduction

For faithful expression of the genetic code, the ribosome must initiate translation only at the start codon. How the ribosome avoids aberrant initiation at other AUG codons is not well understood, although pioneering work suggests certain mRNA sequence elements like the Shine-Dalgarno site are positive determinants for translation initiation at the start codon. In addition, it is known that ribosomes initiate more efficiently if the region surrounding the start codon (the translation initiation region, or TIR) has low secondary structure content (1–3). More complex analyses of TIRs have built large libraries of mRNA mutants and created predictive models of translation initiation based upon these sequence features, which can somewhat predict the initiation level of mRNAs (4). Importantly, such analyses have been performed almost exclusively in *E. coli*, and the importance of the SD has been assumed to be universal among bacteria as the complimentary anti-SD in rRNA is universally conserved across bacteria. Genome-wide analysis of the TIRs of mRNAs in other bacteria has revealed that SD sites are absent in the TIR of many species (5–7), suggesting SD sites are not required for initiation in many species. Indeed, it has been found that some bacteria contain large fractions of leaderless mRNAs which do not even contain a 5’ UTR, making it impossible to make SD-antiSD base pairing interactions (6). Further genome wide analysis revealed that SD sites are also prevalent within the coding sequence where they may induce ribosome pausing and are predicted to be non-functional for initiation (8–10), although neither has been thoroughly tested. The bacterium *C. crescentus* is the fastest growing species that is predominantly nonSD in translation initiation and it also has a well annotated transcriptome (11,12), making it an ideal model to probe how mRNA sequence features program translation initiation in TIRs as opposed to AUGs in the CDS (called elongator AUG regions henceforth) in predominantly nonSD species.

Therefore, we comprehensively examined the roles of SD sites and mRNA secondary structure in the leadered TIRs and elongator AUG regions (EARs) of *C. crescentus*. Using an *in vivo* translation initiation reporter assay, we find that as predicted, ribosomes have a strong preference for initiation on TIRs as compared to EARs. Upon global mRNA sequence analysis, we found more EARs containing SD-AUG pairs than TIRs, but mRNA secondary structure content was lower for TIRs as compared to EARs. We systematically tested the effects of secondary structure content and the presence or absence of SD sites in a combination of TIR and EAR mutants. As expected, the TIR mutants showed that lower secondary structure content and the presence of an SD site can stimulate efficient TIR initiation. EAR mutants that lower mRNA secondary structure content, making the EAR more accessible to ribosomes, stimulates initiation regardless of SD content. Interestingly, EAR mutants with low secondary structure content showed additional stimulation when combined with SD sites; however, EARs containing SD sites within stable secondary structures showed no benefit. Together, this suggests that the lower secondary structure content observed in TIRs as compared to EARs is likely the major determinant of start codon selection and that SD sites tune translation initiation efficiency if accessible to base pair with ribosomes.

## Materials and methods

### Cell growth and media

#### *E. coli* culture

For cloning, plasmids with the reporter gene were transformed in *E. coli* top10 competent cells using the heat shock method for 50-55 secs at 42°C. Luria-Bertani (LB) liquid media was used for outgrowth and the colonies were plated on LB/kanamycin (50 μg/mL) agar plates.

For miniprep, the *E. coli* cultures were inoculated overnight(O/N) in liquid LB/kanamycin (30 μg/mL).

#### *C. crescentus* culture

For cloning, plasmids were transformed in NA1000 *C. crescentus* cells after sequence verification using electroporation. The *C. crescentus* NA1000 cells were grown in Peptone Yeast Extract (PYE) liquid medium. After transformation, for the outgrowth liquid PYE medium was used (2mL) and then plated on PYE/kanamycin (25 μg/mL) agar plates. For imaging, the *C. crescentus* culture were grown O/N at different dilutions in liquid PYE/kanamycin (5 μg/mL). The next day, the cultures growing in log phase were diluted and induced in liquid PYE/kanamycin (5 μg/mL) with xylose (final concentration of 0.2%) such that the optical density (OD) was around 0.05 to 0.1.

### Design and generation of translation reporters

#### Oligo and plasmid design

For the design and generation of reporter assay, a plasmid with a reporter gene (yellow fluorescent protein (YFP)), under the control of an inducible xylose promoter was used. The pBYFPC-2 plasmid containing the kanamycin resistant gene was originally generated from (13). A list of sequences and oligos used for generating plasmids with different TIR and EARs driving translation of YFP is attached as a supplementary table (Table 1).

#### Inverse PCR mutagenesis and ligation

The 5□UTR region and start codon of the YFP reporter protein was replaced with other TIR sequences. This was done by inverse PCR, in which the leaderless TIR was attached to the reverse primer as an overhang. Initial denaturation was done at 98°C for 5 mins, followed by 30 cycles of denaturation at 98°C for 10 secs, annealing at 60°C for 10 secs and extension at 72°C for 7 mins and 20 secs. After 30 cycles, final extension was done at 72°C for 5 mins. The polymerase used was Phusion (Thermoscientific 2 U/μL). The PCR product was DPNI treated to cut the template DNA using DPNI enzyme (Thermoscientific 10 U/μL). The DPNI treated sample was then purified using Thermo fisher GeneJET PCR Purification kit. The purified sample (50 ng) was used for blunt end ligation using T4 DNA Ligase (Thermoscientific 1 WeissU/μL).

#### Transformation in *E. coli* cells

5 μL of the ligation reaction was added to 50 μL of E. coli top10 competent cells. The mixture was incubated in ice for 30 mins. The transformation mixture was heat shocked for 50-55 secs at 42°C, then immediately kept on ice for 5 mins, after which 750 μL of LB liquid medium was added to the cells for outgrowth and kept for incubation at 37°C for 1 hr at 200 rpm. After this, 200-250 μL of the culture was plated on LB/kanamycin (50 μg/mL) agar plates.

#### Colony screening and sequence verification

The colonies grown on LB/kanamycin plates were screened by colony PCR for the presence of the mutant TIR insert. The cloning results in the replacement of the larger 5□UTR region of YFP with a smaller region containing a leaderless TIR, which are easily distinguished on an analytical gel. The forward and reverse primer used for the screening result in a product of approximately 180 base pairs, whereas the original fragment amplified with the same oligos is 245 base pairs. The forward oligo used was pxyl-for: cccacatgttagcgctaccaagtgc and reverse oligo is eGYC1: gtttacgtcgccgtccagctcgac. Upon verification, a small aliquot (4 μL) of the colony saved in Taq polymerase buffer was inoculated in 5 mL of liquid LB/kanamycin (30 μg/mL) and incubated overnight at 37°C at 200 rpm. The next day, the culture was miniprepped using Thermo fisher GeneJET Plasmid Miniprep kit. The concentration of DNA in the miniprepped samples were measured using Nanodrop 2000C from Thermoscientific. DNA samples were sent to Genewiz for Sanger sequencing to verify the correct insert DNA sequences using the DNA primer eGYC1: gtttacgtcgccgtccagctcgac(13).

#### Transformation in *C. crescentus* NA1000 cells

After the sequences were verified, the plasmids were transformed into *C. crescentus* NA1000 cells. For transformation, the NA1000 cells were grown overnight at 28°C in PYE liquid medium at 200rpm. The next day, 5 mL of cells were harvested for each transformation, centrifuged, and washed three times with autoclaved milliQ water. Then, 1 μL of sequence verified plasmid DNA was mixed with the cells and electroporated using the Bio-Rad Micropulser (program Ec1 set at voltage of 1.8 kV). The electroporated cells were immediately inoculated into 2 mL of PYE for 3 hours at 28°C at 200rpm. Then, 10-20 μL of culture was plated on PYE/ kanamycin agar plates. Kanamycin-resistant colonies were grown in PYE/kanamycin medium overnight and then stored as a freezer stock in the −80°C freezer.

#### Cellular assay of translation reporters

*C. crescentus* cells harboring reporter plasmids were serially diluted and grown overnight in liquid PYE/kanamycin medium (5 μg/mL). The next day, cells in the log phase were diluted with fresh liquid PYE/kanamycin (5 μg/mL) to have an optical density (OD) of 0.05-0.1. The inducer (xylose) was then added in the medium such that the final concentration of xylose was 0.2% and the cells were grown for 6 hours at 28°C at 200 rpm. After this, 2-5 μL of the cultures were spotted on M2G 1.5% agarose pads on a glass slide. After the spots soaked into the pad, a coverslip was placed on the pads and the YFP level was measured by fluorescence microscopy using a Nikon eclipse NI-E with CoolSNAP MYO-CCD camera and 100x Oil CFI Plan Fluor (Nikon) objective. Images were captured using Nikon elements software with a YFP filter cube with exposure times of 30 ms for phase-contrast images and 300 ms for YFP images respectively. For EAR experiments, 700 ms exposure for YFP was utilized due to the lower YFP levels. The images were then analyzed using a plugin of software ImageJ(14) called MicrobeJ(15) to calculate the YFP/Cell values.

### Computational predictions of start codon and elongator AUG region accessibility

#### Retrieving transcript sequences

All translation initiation region sequences were retrieved from transcription start site and translation start site data available from RNA-seq and ribosome profiling, respectively (11,12,16), using the *C. crescentus* NA1000 genome sequence (17) using the in-built python scripts.

For elongator AUG regions, all the AUGs within the mRNA sequence were scanned (both in-frame and out of frame AUGs) and then 50 bases were retrieved (−25 from elongator AUG and +25 from elongator AUG, including the AUG).

### ΔG_unfold_ computation

Start codon accessibility was computed similar to (18) as ΔG_mRNA_ - *ΔGInit* using the ΔGunfoldleaderless package (19). This was done in the following three steps:

1. Calculation of ΔG_mRNA_: The minimum free energy (MFE) labelled as ΔG_mRNA_ was calculated using the RNAfold web server of the Vienna RNA websuite (20) at 28°C by inputting all the TIR sequences in a text file (no fasta format required) using the command line function (RNAfold --temp 28 <input_sequences.txt >output.txt). The output file was in the default RNAfold format with each new sequence on one line followed by dotbracket notation (Vienna format) in the next line. The file format was then changed to fasta format so that each sequence and its dot-bracket notation could be inputted to RNAstructure (21) to generate ct files for each sequence. Using the ct file data, all the base pair indexes for each sequence were retrieved and stored in a list assigned for that sequence. Also, the Vienna format of all sequences was extracted from the RNAfold output file and printed on each line of a new text file in the same order as the order of sequences.
2. Calculation of ΔG_Init_: Using these pairing indexes data and original Vienna format, the sequences were constrained such that all the base pairs in the ribosome binding site (RBS) (from up to 12 bases upstream of the start codon to 13 bases downstream of the start codon) were broken and forced to be single stranded while the secondary structures outside the RBS were unchanged. If the 5’UTR length was more than or equal to 25 bases, then the RBS was selected from −12 to +13 bases (25 bases). If the 5’UTR length was less than 25, then the RBS comprised of entire 5’UTR to +13 bases. This was done to ensure that the AUG position relative to the ribosome remains unchanged for different sequences having varying 5’UTR lengths. To calculate the MFE of all the sequences with the constraints, a new file having of the constraint for each sequence is followed by the sequence itself was then input in the RNAfold program (20). This MFE was labelled as ΔG_init_.
3. Calculation of ΔG_unfold_: Lastly, ΔG_unfold_ was calculated by subtracting ΔG_mRNA_ (mfe of mRNA in native state) from *ΔGInit* (mfe of mRNA after ribosome binding) (eq 1. ΔG_unfold_ – ΔG_mRNA_- ΔG_Init_).

## Results

### Translation initiation reporter assay

For proper gene expression ribosomes must be able to correctly distinguish the start codon AUG, which occurs in the translation initiation region (TIR), from other elongator AUGs present in the mRNA (referred to as elongator AUG regions (EARs)). To study what determines the preference for TIRs over EARs, we utilized a translation initiation reporter assay where the start codon for the YFP gene is replaced with the TIR or EAR of mRNAs from the *C. crescentus* transcriptome, forcing the YFP reporter to initiate with either a TIR or EAR (Fig 1A). To validate this *in vivo* translation initiation reporter system, we cloned several leadered translation initiation regions (TIRs) and elongator AUG regions (EARs) from the transcriptome of *C. crescentus* into the pBXYFPC-2 plasmid replacing the 5’ TIR of YFP with one of 6 TIRs or 19 EARs. Upon induction with xylose, the constructs with plasmids having TIR sequences were initiated more efficiently than those with EAR sequences, which were typically very close to background fluorescence levels (Fig 1B). Three of the TIRs contained Shine-Dalgarno (SD) sites, while the other three contained mRNA leaders lacking SD sites. Translation of the two SD TIRs was high, while the other TIRs were translated at moderate levels, in line with prior evidence that SD sites stimulate translation initiation (4,22,23). Interestingly, two EAR regions (from CCNA_00326 and CCNA_00499 CDSs) did show detectable levels of translation initiation (Fig 1C).

**Figure 1.**
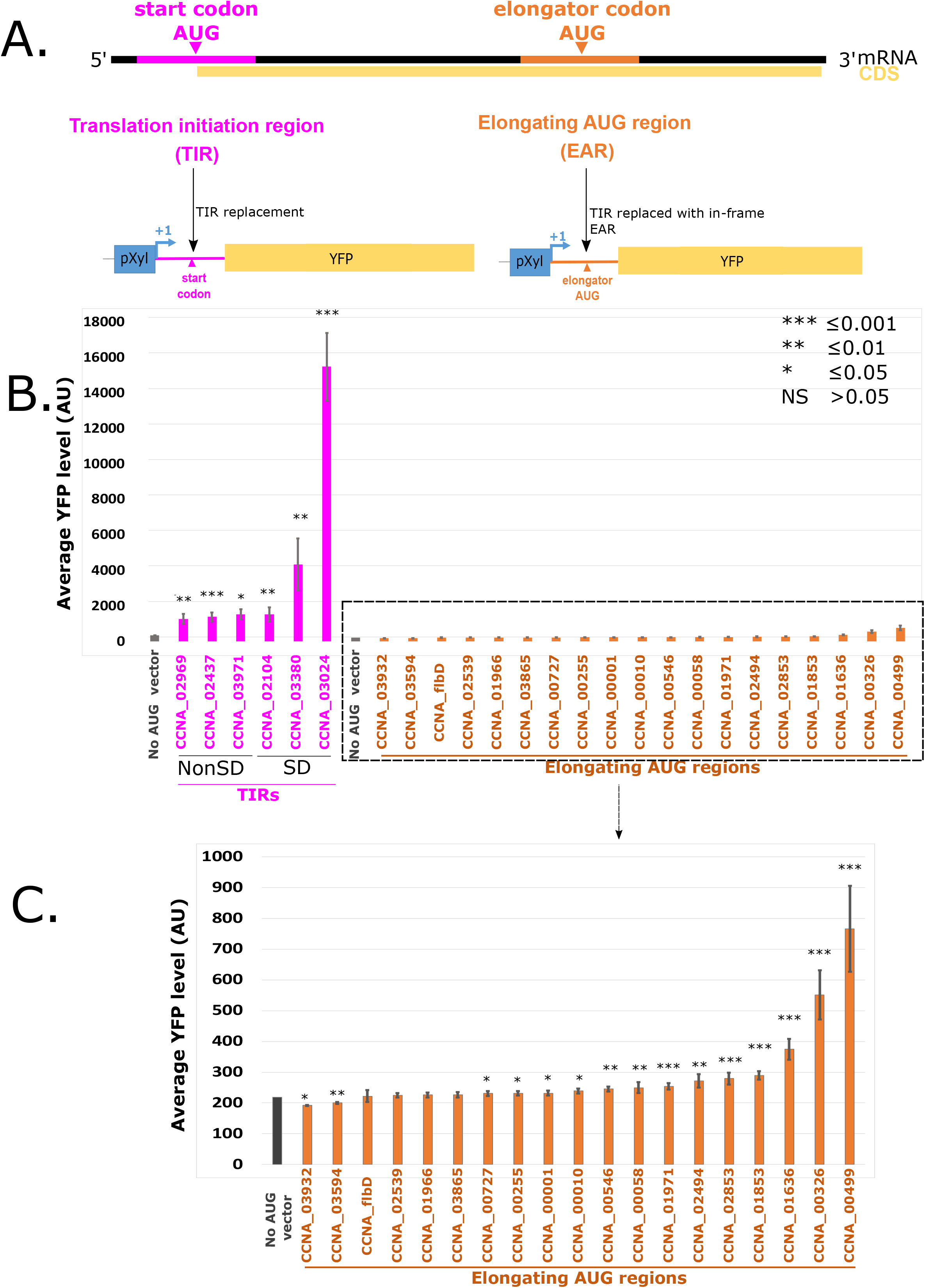
An *in vivo* translation initiation reporter assay shows greater initiation in translation initiation regions (TIRs) as compared to elongator AUG regions (EARs). A.) A graphical representation showing the mRNA with the translation initiation region (TIR) highlighted in pink and an elongator AUG region (EAR) highlighted in orange. Both the TIR and EAR are cloned into a translation initiation reporter plasmid downstream of the pXyl TSS, replacing the start codon of yellow fluorescent protein (YFP), so that translation initiation must occur at the transplanted TIR or EAR to yield YFP fluorescence. B.) Bar chart showing the average YFP intensity of TIRs in pink, EARs in orange, and a vector lacking an AUG start codon in grey. TIRs containing SD sites are indicated. T-test with unequal variances was done to compare all constructs to the no AUG vector with *** representing a p-value of ≤0.001, ** representing a p-value of ≤0.01, * representing a p-value of ≤0.05. C. Zoomed in view of the low reporter levels of EARs from Fig 1B.

### TIRs are predicted to be more accessible than EARs

To determine what mRNA features might promote translation initiation by TIRs and inhibit initiation of EARs, we analyzed the ΔG_unfold_ of all TIRs and EARs in the *C. crescentus* transcriptome. Prior studies have indicated that ribosome accessibility plays a key role in translation initiation (1,2). Accessibility to the ribosome can be approximated by ΔG_unfold_(19), which approximates the free energy required to disrupt the mRNA secondary structure at the TIR to allow ribosomes to initiate (Fig 2A). When applied to all the TIRs and EARs in the genome, we saw that TIRs showed a lower ΔG_unfold_ on average, with little difference between in frame EARs or out of frame EARs, suggesting that TIRs are more accessible to ribosomes (Fig 2C). Across the transcriptome, there are 2803 leadered TIR regions, while there are 36,291 EAR regions, making EARs far more abundant than TIRs. As SD sites are known to stimulate initiation via base pairing of the mRNA to the anti-SD in the 16S rRNA (Fig 1B), we analyzed the presence of SD sites across all TIR and EAR regions in the transcriptome. SD sites were defined conservatively as 4 continuous bp with the core of the anti-SD occurring within 20nt upstream of the start codon. Despite the textbook importance of SD sites in the TIR, we observed only 1335 SD-AUG pairs in TIRs, while we found 6850 in EARs (4399 in frame, 2451 out of frame). Even though there are more SD-AUG pairs in EARs than TIRs, the probability of finding an SD-AUG pair is higher in TIRs than EARs (Fig 2B). As the *C. crescentus* genome is very GC rich (67.3% GC), the probability of finding an SD site in a random sequence is 28.5%; TIRs have more SD sites than expected by random chance, while EARs have less SD sites than random chance (Fig 3B).

**Figure 2.**
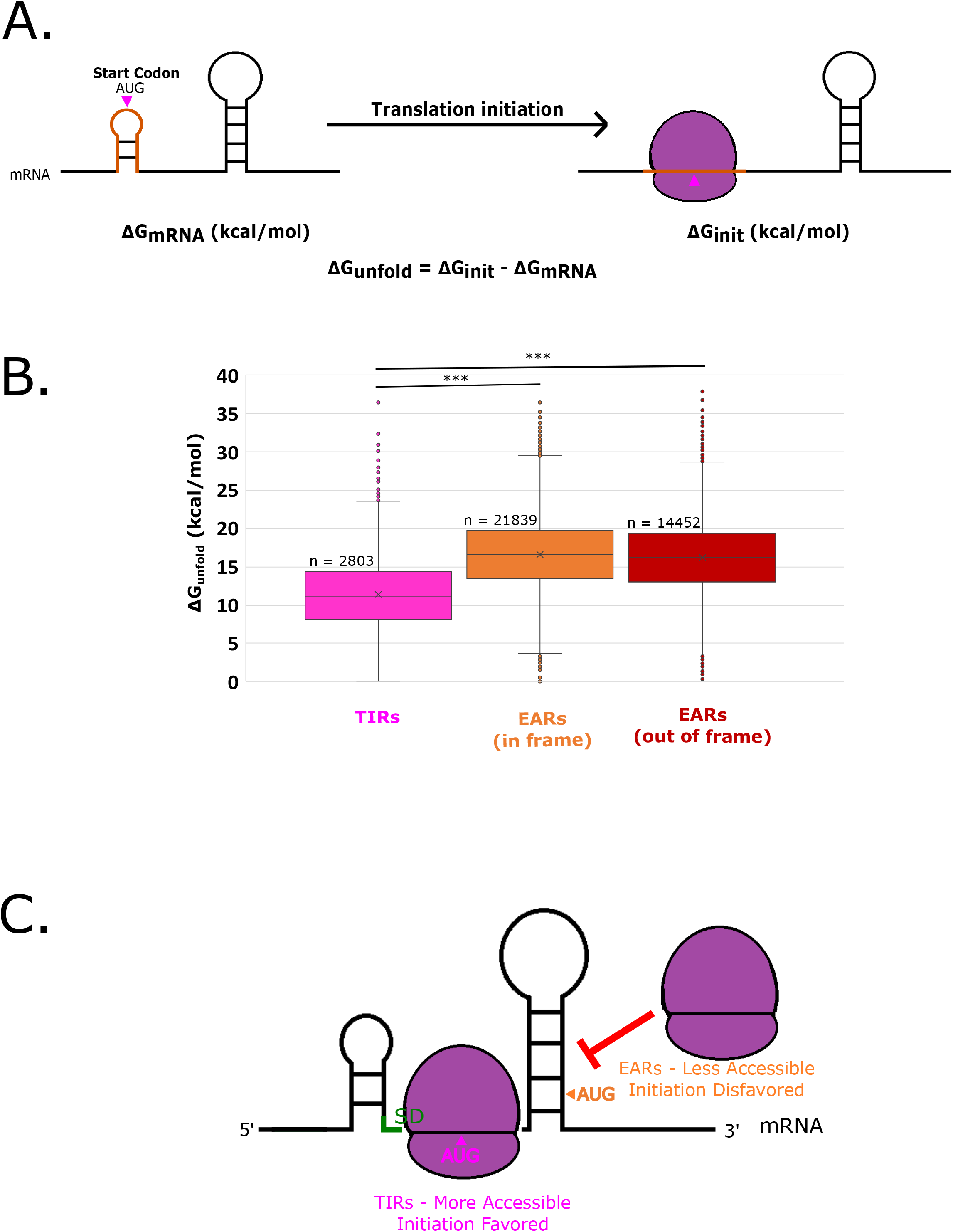
EARs are less accessible than TIRs across the *C. crescentus* transcriptome. A.) mRNA accessibility can be estimated by the calculation of ΔG_unfold_. The predicted mRNA minimum free energy (ΔG_mRNA_) is represented on the left. The orange translation initiation region indicates a ribosome footprint surrounding the start codon (pink). The image on the right represents the mRNA upon initiation (ΔG_init_), where the orange initiation region is unfolded by the ribosome. The ΔG_unfold_ represents the amount of energy required to unfold the translation initiation region of the mRNA. B.) Box-whisker plot showing the distribution of ΔG_unfold_ across all mapped TIRs and EARs in the *C. crescentus* transcriptome (19). Pink box-whisker represents TIRs, orange represents in-frame EARs, and red represents out-of-frame EARs. *** indicates a p-value of ≤0.001 for 2-tailed T-test with unequal variance. C.) A graphical representation showing that TIRs are generally more accessible, facilitating initiation, whereas EARs are less accessible, blocking ribosome access.

**Figure 3.**
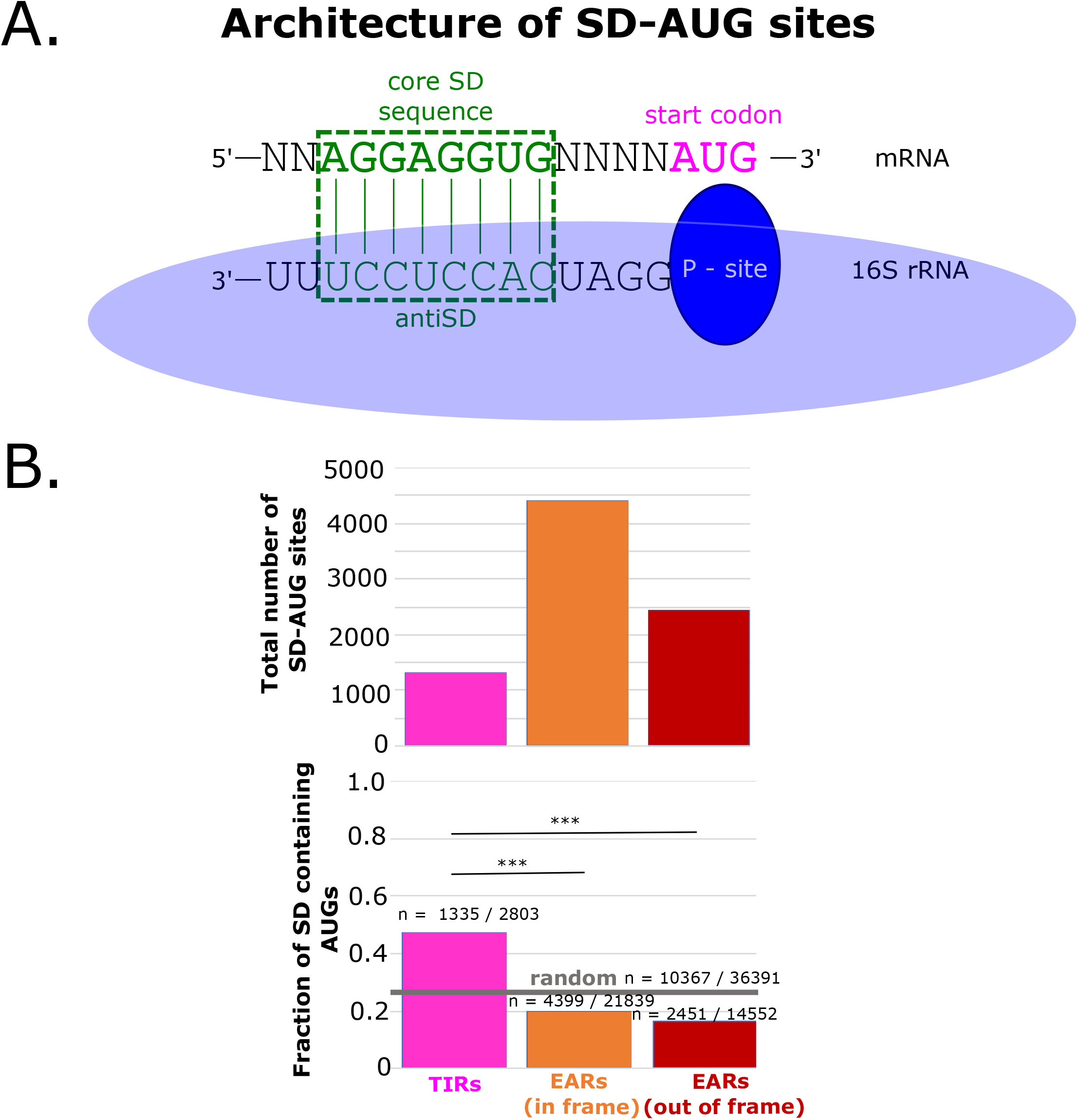
Shine-Dalgarno-AUG pairs are more abundant in EARs than TIRs. A.) A graphical representation of the optimal alignment of the core SD sequence “AGGAGGUG” in the mRNA in green with the anti-SD sequence in the rRNA below. The base pairing is highlighted in the green dotted box. B.) The abundance of SD-AUG pairs across TIRs and EARs. The total number of SD-AUG pairs TIRs and EARs is on the top. The pink bar represents TIRs, the orange bar represents in-frame EARs and the red bar represents out-of-frame EARs. Below is the fraction of SD enrichment in TIRs and EARs. The random probability of SD enrichment is shown as grey horizontal line (estimated from 36391 random sequences of *C. crescentus’* 67% genomic GC%). *** represents p-value of ≤0.001 calculated from 2-sample Z-test.

Interestingly, SD sites in TIRs occur at a higher frequency between 15 and 10nt upstream of the AUG (24), while SD sites in EARs occurs more often with non-optimal spacing for initiation (Fig 4A). To test whether the position of the SD site affects initiation in *C. crescentus*, we generated TIR reporters with an altered position of the SD site (Fig 4A,B). The TIR reporters were designed with a poly-A 5’ UTR, which minimizes the chances of inhibitory RNA secondary structures forming, to minimize potential effects of mRNA structure content on SD spacing. The translation initiation reporters with altered SD position showed that SD sites located within the optimal range (10-15nt upstream) led to ~4-6 fold higher translation initiation efficiency as compared to a nonSD control TIR (Fig 4B), while those located close to the start codon (7 or 9nt upstream) or far upstream (17nt) showed insignificant changes in translation initiation efficiency compared to the nonSD control. Taken together, this suggests there is likely some positive selection for optimally spaced SD sites in TIRs, and a slight negative selection against optimally spaced SD sites in EARs. In addition, EARs tend to have more SD sites located outside the optimal region which likely will not stimulate initiation. Overall, SD-AUG pairs are prevalent in both TIRs and EARs and have modest effects on translation initiation but cannot fully explain the strong preference for initiation at TIRs over EARs (Fig 1).

**Figure 4.**
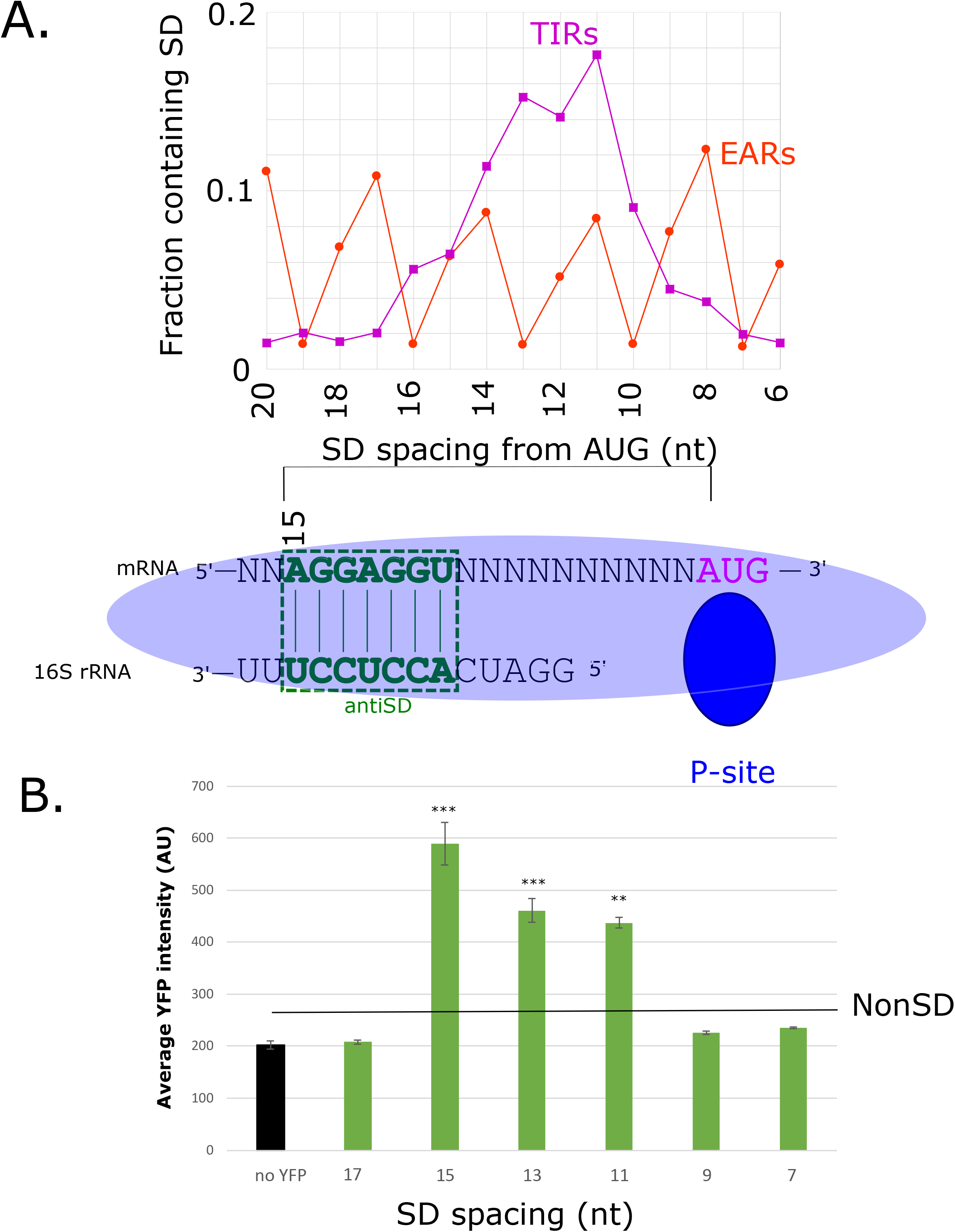
Strong, optimally spaced SD-sites boost translation efficiency of a TIR. A.) Distribution of SD site spacing in TIRs (dark magenta) and EARs (orange) across the *C. crescentus* transcriptome. Aligned spacing is calculated from the 5’ end of the core SD site after alignment to the anti-SD as shown below. B.) Bar chart showing the average YFP production in the control (empty vector) compared to plasmids with a poly-A 5’ UTR and SD sites spaced relative to the AUG as indicated below. The poly-A 5’ UTR was chosen as it limits the base pairing of the SD site with other bases in the TIR. A T-test with unequal variances was done to compare all constructs with nonSD control with *** representing a p-value of ≤0.001, ** representing a p-value of ≤0.01, * representing a p-value of ≤0.05.

### mRNA accessibility is a major determinant of start codon selection

As both mRNA accessibility and the presence of an SD site can promote translation initiation, we designed a series of EAR mutants where we combinatorially altered mRNA secondary structure, the presence of an SD site, or both with the goal of converting these EAR sites into functional TIRs (Fig 5). First, we examined the EARs in our translation initiation reporter system and found that approximately half have SD sites upstream of the AUG codon. We calculated the ΔG_unfold_ and found that EARs with higher ΔG_unfold_ were more poorly translated than those having a lower ΔG_unfold_ (Fig 5A, 488.63 average high accessibility, 247.18 average low accessibility). We did observe a slight increase in initiation reporter levels in the SD containing EARs (229.08 avg nonSD, 320.85 avg SD); however, it was not statistically significant based on a Mann Whitney U test used to compare skewed distributions (Fig 5B). We then introduced combined mutations in the 5’ UTRs of these EARs mutants that would alter the presence of an SD site, ribosome accessibility, or both simultaneously. Six of these EARs were nonSD in the wild type and six contained an SD site in the wild type. Across the nonSD EARs, we observed a detectable increase in initiation levels in EARs that were highly accessible (average YFP level highly accessible 302.25, moderate 242.31, low accessibility 225.31, High vs low p-value .01, high vs moderate p-value 0.04, Fig 5C). We observed that SD sites did not significantly stimulate translation if the EAR region has low accessibility (p-value=0.154). At moderate levels of mRNA accessibility, we observed that some EARs with SD sites were initiated at higher efficiency, with a p-value of 0.055. However, in EAR mutants that were both highly accessible, SD site containing versions showed a strong stimulation of translation initiation into the range of the natural TIR reporters (Fig 5C, Fig 1B). This suggests that EARs are strongly prevented from initiating due to their general lack of accessibility to ribosomes, and that SD sites only stimulate initiation if the EAR region is accessible to ribosomes.

**Figure 5.**
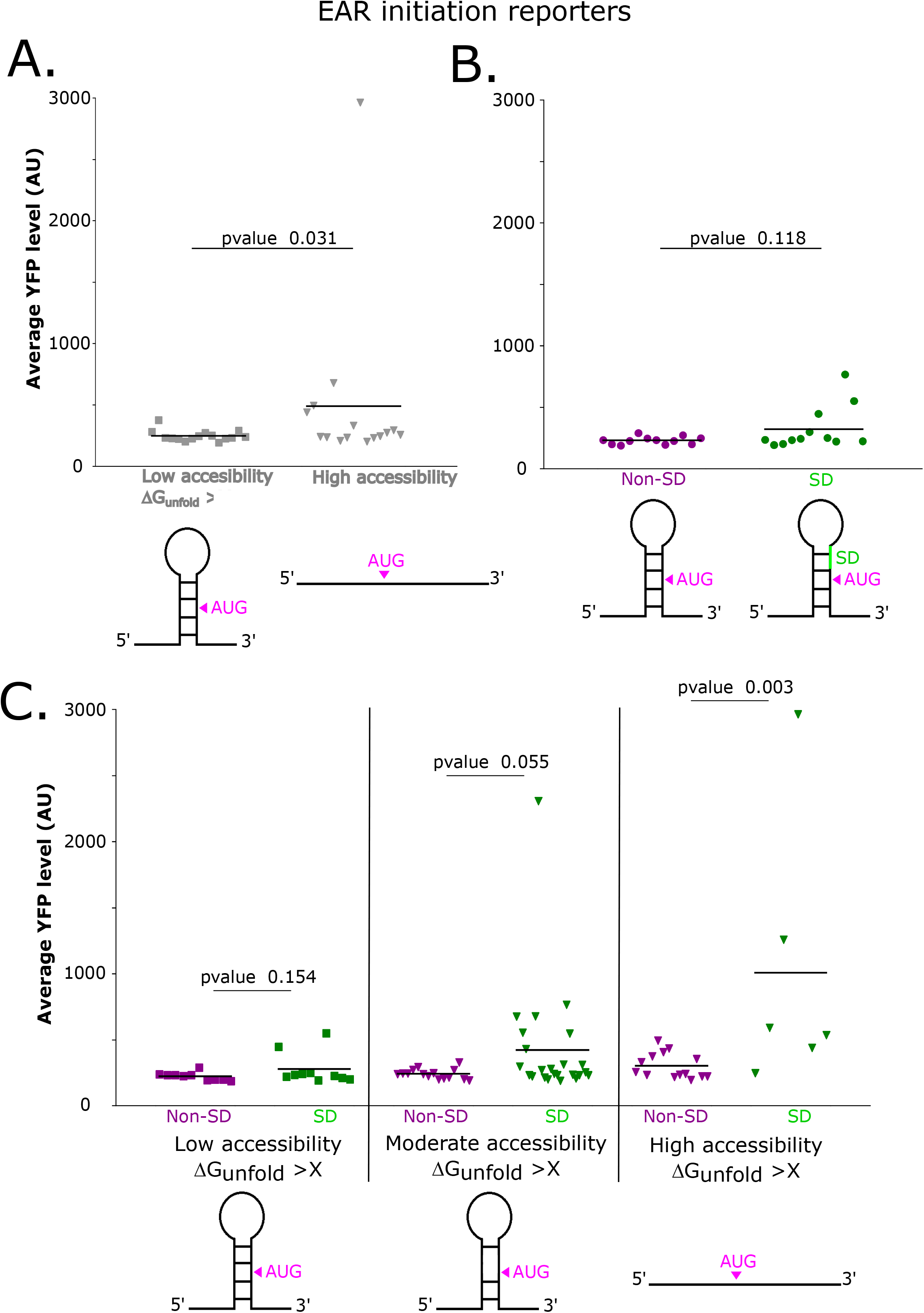
Low EAR accessibility prevents stimulation of initiation by SD sites. A.) Distribution plot showing average YFP levels of non-accessible wild-type EARs represented by square data points vs accessible mutant EARs represented by inverted triangle data points. EAR mutants contain point mutations in the region upstream of the AUG that reduce potential base-pairing in the EAR region. Each point represents a single EAR reporter construct’s *in vivo* YFP level (Table 2). A Mann-Whitney U test was calculated between low accessibility and high accessible constructs to assess significance. B.) Distribution plot showing average YFP level of non-SD EARs represented by dark magenta data points vs SD EARs represented by green data points. A Mann-Whitney U test between non-SD and SD EAR initiation reporters was used to assess significance. C.) Distribution plot showing average YFP of non-SD and SD TIRs with different degrees of accessibility. Dark magenta data points represent non-SD TIRs and green data points represents SD TIRs. Squared data points represent low accessibility TIRs and inverted triangle data points represent moderately accessible and highly accessible TIRs. A Mann-Whitney U test was performed for non-SD and SD pairs in each accessibility category to compare skewed distributions for significance.

To further investigate how AUG accessibility and SD content coordinate start codon selection we analyzed their relationship as compared to translation efficiency across TIRs in the *C. crescentus* transcriptome (Fig 6A). Here we observed that the proportion of SD sites in EARs was highest if the EAR had low accessibility, while the proportion of SD sites dropped in moderate or highly accessible EARs. Interestingly, in TIRs we observed a similar decrease in the proportion of SD sites in highly accessible TIRs. To investigate whether natural TIRs are stimulated by accessibility and SD content, we examined translation efficiency (TE) as determined by ribosome profiling (Fig 6B) (11). For this analysis, TIRs from only monocistronic or the first genes in operons were used to avoid the effects of translational coupling on the TE measurements. Across non-SD mRNAs, we observed that as accessibility increases, average TE also increases from 0.75 to 1.0. A similar trend occurs for SD mRNAs, where the average TE increases from 0.83 to 1.6. In addition, we observe that as the TIRs become more accessible, the SD mRNAs have increasing levels of translation efficiency, suggesting that highly accessible SDs also stimulate translation initiation efficiency of natural mRNAs. In the low accessibility TIRs, we observe a small difference in average TE between nonSD (0.75) as compared to SD (0.82); however, the p-value (0.09) only gives a low confidence interval. As we compared the less structured TIRs in the moderate or high accessibility bins, we observed larger average TE values and higher confidence intervals between nonSD and SD mRNAs. The average TE observed for highly accessible mRNAs was 1.0 in nonSD mRNA and 1.6 in SD mRNAs (p-value = 8.2E-4). This supports the conclusion that SD sites do not have significant effects on translation initiation if they are blocked by inhibitory secondary structures, but they can positively influence translation efficiency if the SD sites are accessible to base pair with the rRNA.

**Figure 6.**
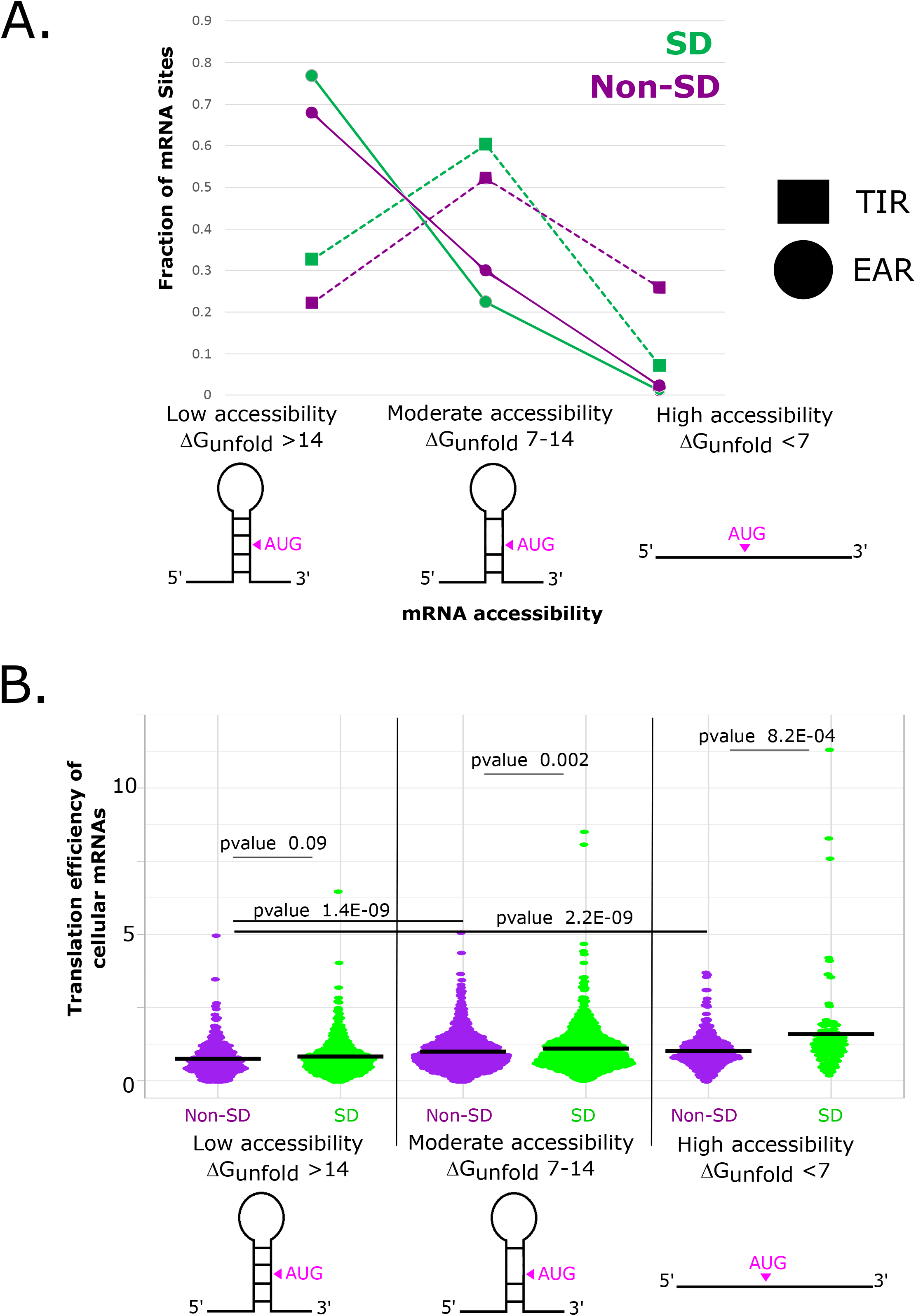
TIRs with accessible SD-AUG pairs have higher translation efficiency in natural mRNAs. A.) Plot showing the distribution of SDs in TIRs and EARs classified with respect to their accessibility. NonSD TIRs/EARs are shown in dark magenta and SD TIRs/EARs are shown in green. B.) Translation efficiency values of *C. crescentus* mRNAs across six categories of accessibility and SD site prevalence as measured by ribosome profiling (11). Dark magenta data points represent non-SD TIRs and green data points represents SD TIRs. Squared data points represent low accessibility TIRs and inverted triangle data points represent moderate accessibility and high accessibility TIRs. P-value of Mann-Whitney U test calculated for non-SD and SD pairs for each accessibility category were used to compare skewed distributions for significance.

## Discussion

### AUG accessibility is a major determinant of start codon selection, while the SD can tune efficiency of initiation

In line with previous observations in other organisms (1–3,7), we find that translation initiation on leadered mRNAs is strongly impacted by the ΔG_unfold_ in *C. crescentus* (Fig 7). A similar observation was also observed in leaderless mRNAs in *C. crescentus* (25), suggesting that a low ΔG_unfold_ may be a universal requirement for initiation regardless of initiation mechanism. Indeed, the large difference in ΔG_unfold_ observed in TIRs and EARs appears to be the major determinant for ribosomes to accurately select the start codon. Despite the textbook view that SD sites direct start codon selection, a larger abundance of SD sites appear in EARs as compared to TIRs (Fig 3B), with those in EARs generally being nonfunctional, most likely because they are blocked from 16S rRNA base-paring due to the higher mRNA secondary structure content present in EARs (Fig 6). Indeed, SD sites in EARs also have a higher propensity to be non-optimally spaced, which reduces their ability to stimulate initiation (Fig 4). SD sites in TIRs promote more efficient initiation in *C. crescentus* (Figs 1,4), suggesting that while the SD site is not required for initiation, it acts to tune initiation efficiency. Interestingly, recent ribosome profiling experiments using *E. coli* ribosomes with a mutated anti-SD showed that they still initiate at the original start codon, albeit with altered translation initiation efficiency (26). This suggests that even in *E. coli*, where the SD sequence was originally proposed to be the major determinant for translation initiation, it is instead used to tune translation initiation efficiency. The lack of secondary structure appears to be a universal feature of bacterial TIRs (3,7), while SD site content in the TIR is highly variable across many clades (5). In the Bacteroidetes clade, where SD site content in TIRs is very low (5), a cryo-EM structure of a 70S initiation complex revealed that its anti-SD is prevented from base pairing with mRNA by ribosomal proteins bS21 and bS18 (27). This suggests that the SD site is not used universally to tune initiation efficiency in all bacteria even though the anti-SD rRNA sequence is universally conserved. Indeed, many organisms are known to initiate translation with leaderless mRNAs that are completely devoid of a 5’ UTR and lack SD sites (6,25). Intriguingly, the frequency of SD sites in the CDS was found to be inversely correlated with growth rate across bacteria (9), suggesting that SD sites in EARs may be more detrimental in fast growing species. Moving forward, it will be critical to determine the functional importance of TIR accessibility and SD sites to translation initiation across diverse species of bacteria.

**Figure 7.**
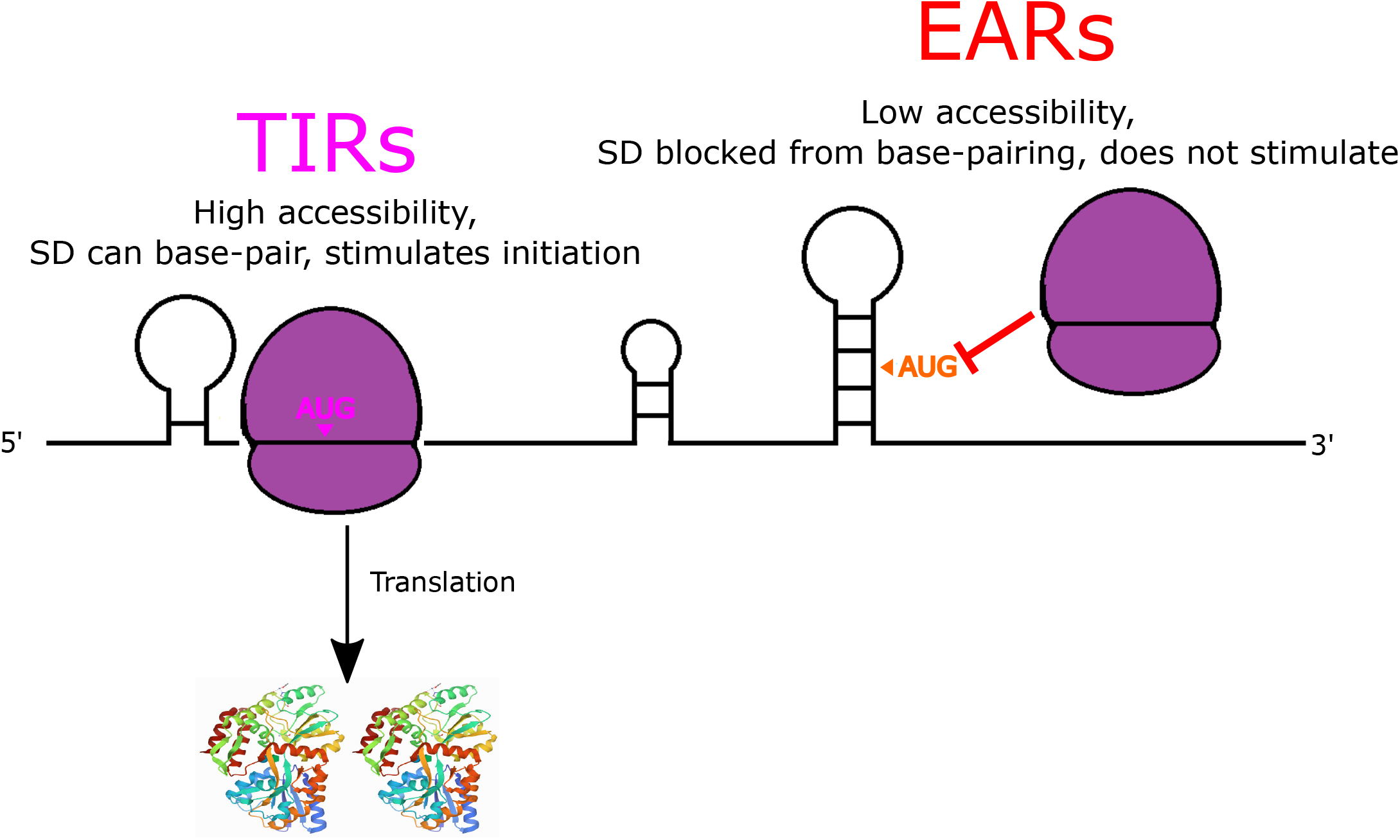
AUG accessibility dictates start codon selection while the SD can boost initiation efficiency. Cartoon showing that TIRs are more accessible than EARs, promoting TIR initiation (dark magenta) and preventing initiation on EARs (red orange). SD-AUG pairs are abundant in both TIRs and EARs, but only increase initiation efficiency in TIRs where the mRNA is highly accessible.

## Acknowledgement

This work was funded by grant R35GM124733 by NIGMS. The authors thank Adam Hockenberry for in depth discussions and Erin Schrader for critical reading of the manuscript.

## Author Contributions

JMS designed study, obtained funding, and oversaw experiments and data analysis. MHB, AsG, and AmG cloned plasmids, generated strains, performed experiments, and analyzed data. AsG, MHB, and JMS wrote the paper.

## Supporting information

Tables and Figure legends

Table 1: List of oligos used for cloning translation initiation reporters.

**XLSX**

Table 2: mRNA features and translation levels of all TIR and EAR initiation reporters.

**XLSX**

## Notes

### Competing Interest Statement

The authors have declared no competing interest.

## References

1. de Smit, M.H. and van Duin, J. (1990) Secondary structure of the ribosome binding site determines translational efficiency: a quantitative analysis. Proceedings of the National Academy of Sciences of the United States of America, 87, 7668–7672.

2. Mustoe, A.M., Busan, S., Rice, G.M., Hajdin, C.E., Peterson, B.K., Ruda, V.M., Kubica, N., Nutiu, R., Baryza, J.L. and Weeks, K.M. (2018) Pervasive Regulatory Functions of mRNA Structure Revealed by High-Resolution SHAPE Probing. Cell, 173, 181–195.e118.

3. Gu, W., Zhou, T. and Wilke, C.O. (2010) A universal trend of reduced mRNA stability near the translation-initiation site in prokaryotes and eukaryotes. PLoS computational biology, 6, e1000664.

4. Salis, H.M. (2011) The ribosome binding site calculator. Methods Enzymol, 498, 19–42.

5. Chang, B., Halgamuge, S. and Tang, S.-L. (2006) Analysis of SD sequences in completed microbial genomes: non-SD-led genes are as common as SD-led genes. Gene, 373, 90–99.

6. Beck, H.J. and Moll, I. (2018) Leaderless mRNAs in the spotlight: ancient but not outdated! Microbiology spectrum, 6, 6.4. 02.

7. Scharff, L.B., Childs, L., Walther, D. and Bock, R. (2011) Local absence of secondary structure permits translation of mRNAs that lack ribosome-binding sites. PLoS genetics, 7, e1002155.

8. Hockenberry, A.J., Jewett, M.C., Amaral, L.A.N. and Wilke, C.O. (2018) Within-Gene Shine-Dalgarno Sequences Are Not Selected for Function. Molecular biology and evolution, 35, 2487–2498.

9. Yang, C., Hockenberry, A.J., Jewett, M.C. and Amaral, L.A.N. (2016) Depletion of Shine-Dalgarno Sequences Within Bacterial Coding Regions Is Expression Dependent. G3 (Bethesda, Md.), 6, 3467–3474.

10. Li, G.W., Oh, E. and Weissman, J.S. (2012) The anti-Shine-Dalgarno sequence drives translational pausing and codon choice in bacteria. Nature, 484, 538–541.

11. Schrader, J.M., Zhou, B., Li, G.W., Lasker, K., Childers, W.S., Williams, B., Long, T., Crosson, S., McAdams, H.H., Weissman, J.S. et al. (2014) The coding and noncoding architecture of the Caulobacter crescentus genome. PLoS genetics, 10, e1004463.

12. Bharmal, M.-H., Aretakis, J.R. and Schrader, J.M. (2020) An Improved Caulobacter crescentus Operon Annotation Based on Transcriptome Data. Microbiology Resource Announcements, 9, e01025–01020.

13. Thanbichler, M., Iniesta, A.A. and Shapiro, L. (2007) A comprehensive set of plasmids for vanillate-and xylose-inducible gene expression in Caulobacter crescentus. Nucleic Acids Res, 35, e137.

14. Schindelin, J., Arganda-Carreras, I., Frise, E., Kaynig, V., Longair, M., Pietzsch, T., Preibisch, S., Rueden, C., Saalfeld, S., Schmid, B. et al. (2012) Fiji: an open-source platform for biological-image analysis. Nat Methods, 9, 676–682.

15. Ducret, A., Quardokus, E.M. and Brun, Y.V. (2016) MicrobeJ, a tool for high throughput bacterial cell detection and quantitative analysis. Nat Microbiol, 1, 16077.

16. Zhou, B., Schrader, J.M., Kalogeraki, V.S., Abeliuk, E., Dinh, C.B., Pham, J.Q., Cui, Z.Z., Dill, D.L., McAdams, H.H. and Shapiro, L. (2015) The global regulatory architecture of transcription during the Caulobacter cell cycle. PLoS Genet, 11, e1004831.

17. Marks, M.E., Castro-Rojas, C.M., Teiling, C., Du, L., Kapatral, V., Walunas, T.L. and Crosson, S. (2010) The genetic basis of laboratory adaptation in Caulobacter crescentus. J Bacteriol, 192, 3678–3688.

18. Mustoe, A.M., Corley, M., Laederach, A. and Weeks, K.M. (2018) Messenger RNA Structure Regulates Translation Initiation: A Mechanism Exploited from Bacteria to Humans. Biochemistry, 57, 3537–3539.

19. Bharmal, M.-H.M. and Schrader, J.M. (2021) ΔG_unfold_leaderless, a package for high-throughput analysis of translation initiation regions (TIRs) at the transcriptome scale and for leaderless mRNA optimization. bioRxiv, 2021.2008.2027.457836.

20. Gruber, A.R., Lorenz, R., Bernhart, S.H., Neubock, R. and Hofacker, I.L. (2008) The Vienna RNA websuite. Nucleic Acids Res, 36, W70–74.

21. Reuter, J.S. and Mathews, D.H. (2010) RNAstructure: software for RNA secondary structure prediction and analysis. BMC Bioinformatics, 11, 129.

22. Shine, J. and Dalgarno, L. (1974) The 3□-terminal sequence of Escherichia coli 16S ribosomal RNA: complementarity to nonsense triplets and ribosome binding sites. Proceedings of the National Academy of Sciences, 71, 1342–1346.

23. Shine, J. and Dalgarno, L. (1975) Determinant of cistron specificity in bacterial ribosomes. Nature, 254, 34–38.

24. Chen, H., Bjerknes, M., Kumar, R. and Jay, E. (1994) Determination of the optimal aligned spacing between the Shine-Dalgarno sequence and the translation initiation codon of Escherichia coli mRNAs. Nucleic acids research, 22, 4953–4957.

25. Bharmal, M.-H.M., Gega, A. and Schrader, J.M. (2021) A combination of mRNA features influence the efficiency of leaderless mRNA translation initiation. NAR Genomics and Bioinformatics, 3.

26. Saito, K., Green, R. and Buskirk, A.R. (2020) Translational initiation in E. coli occurs at the correct sites genome-wide in the absence of mRNA-rRNA base-pairing. eLife, 9.

27. Jha, V., Roy, B., Jahagirdar, D., McNutt, Z.A., Shatoff, E.A., Boleratz, B.L., Watkins, D.E., Bundschuh, R., Basu, K., Ortega, J. et al. (2021) Structural basis of sequestration of the anti-Shine-Dalgarno sequence in the Bacteroidetes ribosome. Nucleic acids research, 49, 547–567.

